# Mice with a humanized immune system are resilient to transplantation of human microbiota

**DOI:** 10.1101/2021.10.06.463343

**Authors:** Wei Zhou, Kin-hoe Chow, Rory Geyer, Paola Peshkepija, Elizabeth Fleming, Chun Yu, Karolina Palucka, Julia Oh

## Abstract

Human gut microbiota has co-evolved with human, and plays important roles in regulating the development and functioning of the host immune system. To study the human-specific microbiome-immunune interaction in an animal model is challenging as the animal model needs to capture both the human-specific immune functions and the human-specific microbiome composition. Here we combined two widely-used humanization procedures to generate a humanized mouse model (HMA-huCD34) with functional human leukocytes developed from engrafted huCD34+ cells and human fecal microbes introduced through fecal microbiota transplantation, and investigated how the two introduced human components interact. We found that the engrafted human leukocytes are resilient to the transplanted human microbes, while reciprocally the transplanted microbial community in the huCD34 mice was significantly different from mice without a humanized immune system. By tracking the colonization of human fecal Bacteroides strains in the mouse gut, we found that the composition of the strain population changes over time, the trajectory of which depends upon the type of mouse. On the other hand, different from Bacteroides, *Akkermansia muciniphila* exhibited consistent and rapid fixation of a single donor strain in all tested mice, suggesting strong purifying selection common to all mouse types. Our prospect study illustrated the complex interactions between the transplanted microbiome and different host factors, and suggested that the humanized mouse model may not faithfully reproduce the human-specific microbiome-immune interaction.

## Introduction

Mice, because of their genetic similarities to humans, tractability, and short reproductive cycles, have been the most common model system for studying human biology and disease. Nonetheless, some factors of mouse biology, such as aspects of the innate and adaptive immune system and microbiome composition, are considerably discrepant from human ^1,2^. At times, these discrepancies result in limited translation of observations made in the mouse models to humans ^1,3^. For example, numerous drugs for diabetes ^4,5^ and cancer ^6^ have shown different responses or toxicity in mice and human, and many pathogens, such as HIV, are human-specific. To enhance the ability of mouse models to recapitulate human conditions and physiology, “humanized” mouse models have been developed ^3,7,8^, which carry human genes, cells, tissues, organs, and/or human-associated microbes. With these modifications, humanized mouse models have improved translatability to humans in different applications, including host-microbe interactions, drug effects and safety, human-specific diseases, and the human immune system ^3,7–9^.

As an emerging example of humanized mouse models, human microbiota-associated (HMA) mouse models were developed to study the functions of human-associated microbes due to the inextricable impact of the microbiome on health and disease ^7^. HMA mouse models were developed because the mouse gut microbiome differs significantly from the human gut microbiome ^1,10,11^, resulting in substantial discrepancies in interpretation between human conditions studied in mice. For example, the response of the microbiome to dietary interventions or inflammatory bowel disease was found to be significantly different between humans and mice ^1^. HMA mouse models are commonly constructed through fecal microbiota transplantation (FMT) – the inoculation of human fecal microbiota into either mice pretreated with antibiotics or germ-free mice. Conversely, HMA mouse models have been successfully applied to study diseases such as non-alcoholic fatty liver diseases ^12^ and obesity ^13^. Despite HMA mouse models’ popularity, concerns have been raised regarding the ecological variability of the colonized microbiota in the mouse gut environment ^7^. Moreover, mouse models with different genetic backgrounds or non-genetic manipulations (e.g. the engraftment of human cells) differ in their gut physical conditions, nutrient availability, and immune responses, all of which could impose distinct selective pressures on the colonizing microbes ^7^. As a result, colonization dynamics of human microbes could differ across mouse models ^11,14^. We have previously shown that these differences are often manifested at low taxonomic levels: for example, different Bacteroides species were enriched in immunocompetent vs. immunodeficient mice, demonstrating environmental selection at the species level ^11^.

Moreover, even within the same microbial species, different conspecific strains could preferentially colonize certain mouse gut environments but not others depending on interstrainic, interspecific, and environmental or immune selection ^11^. These strain-level differences can be physiologically critical because conspecific strains in the gut microbiome can have diametrically different biological functions in immunomodulation, virulence, and metabolic potential. For example, *Bacteroides fragilis*, a Gram negative anaerobe that commonly colonizes the human gut, has a myriad of strain-specific functions. A subset of *B. fragilis* strains are capable of producing fragilysin – an enterotoxin known to cause diarrheal disease and potentially associated with colorectal cancer ^15^. On the other hand, other *B. fragilis* strains isolated from human fecal samples regulate important functions of dendritic cells (DCs) and macrophages ^16^. Therefore, in addition to species-level profiling which provides community-level information, conspecific strains need to be resolved in the microbiome of HMA mice to correctly interpret mechanistic studies performed using these mouse models.

Another concern for HMA mouse models is that replacement of the microbiota itself might not be adequate to recapitulate host-microbiome interactions that are host-specific. For example, it has been suggested that many aspects of the host immune system, such as the composition of immune cell types and cytokine secretion, could only be educated and regulated by a microbiome that has co-evolved with it ^7,17,18^. This has been strongly supported by a recent study that showed that a common laboratory strain of mice, C57BL/6J, transplanted into wild mice develops a significantly different immune maturation that arguably, better phenocopied human immune responses ^19^. The corollary is that an incompatibility between the host immune system and the microbiome could result in pathological phenotypes. For example, HMA mice models, with a mouse immune system and a human microbiome, can have reduced immune cell numbers and efficiency in clearing infections ^17^. Therefore, HMA mouse models could potentially benefit from additional humanization of the immune system to reconstruct host-specific host-microbiome interactions.

One of the most widely used mouse models with a humanized immune system is the human CD34+ (huCD34) mouse model. HuCD34 mice are constructed by engrafting immunodeficient mice (e.g., NOD.Cg-PrkdcscidIl2rgtm1Wjl/SzJ (NSG) or NSG-SGM3 mice, which express human interleukin-3 (IL-3), human granulocyte/macrophage-colony stimulating factor 2 (GM-CSF), and human stem cell factor (SCF), in the NSG background to support development of human myeloid progenitor and dendritic cells, T and B cells) with human hematopoietic progenitor cells (CD34+ cells), which can diverge and reconstruct the human leukocyte compartments inside the mice ^20^. When engrafted with a human microbiota, HMA huCD34 mice is in theory a suitable model to study questions related to immune-microbiome interactions, ranging from microbiome-modulated immune development to microbiome-dependent efficacy of cancer immunotherapies.

In an HMA huCD34 mouse, the humanized immune environment could shape the FMT microbiota through host-specific immune selection on the colonizing microbes. On the other hand, given the immunomodulatory effect of the microbiome, the FMT microbiota could potentially influence the engrafted human immune cells in huCD34 mice. Timing of FMT administration, or antibiotic treatment could also affect engraftment ^21^. However, to the extent of our knowledge, the colonization dynamics of human microbes in the huCD34 mice, as well as its impact on the engrafted human immune cells, have not been studied in detail. In this prospective study, we created HMA huCD34 mouse models to study microbial transplantation dynamics and the short and long-term impact of the FMT microbiota on the engrafted human immune cells. We found that while the humanized immune environment was associated with a significantly altered post-FMT microbiota, the human microbiota did not significantly alter composition of mature circulating human leukocytes. Additionally, through comparison with other types of HMA mice, we found that different microbial species exhibited distinct strain-level colonization dynamics, revealing different sources of selective forces collectively driving the population structures of foreign microbial colonizers. Finally, we administered an antibiotic pulse and FMT early during huCD34 engraftment and observed few significant long-term differences in human cell engraftment efficiency or immune cell frequencies as a result of no- or human-specific microbial interactions. Our results suggest that on a community scale, human microbiota are, largely, immunogenically equivalent to mouse microbiota in its short term impact on systemic and circulating immunity in the huCD34 system, but that the presence of different human immune compartments in immunodeficient mice could exert selective forces against exogenous colonization.

## Results

### Pre-FMT mouse microbiota of huCD34 and SGM3 mice

Reference-based metagenomic profiling methods have performed poorly on native mouse microbiota to date due to a paucity of mouse-associated microbes in reference databases such as GenBank ^11^. Therefore, we leveraged a recent reported integrated mouse gut metagenome catalog (IMGMC) of metagenomic assembled genomes (MAGs) retrieved from multiple mouse metagenomic whole-genome shotgun sequencing (mWGS) projects ^22^, combined with MAGs reconstructed *de novo* from the mWGS data in this study using a binning method previously described ^11^. From the pre-FMT huCD34 and SGM3 samples, we extracted 53 high-quality MAGs with greater than 90% completeness and lower than 5% contamination (Supplementary Table 1). The 53 high quality MAGs were then combined with 830 high-quality IMGMC MAGs to improve taxonomic representation. Redundant MAGs in the IMGMC catalog that were highly similar to any of the 53 MAGs identified in this study (at least 96.5% average nucleotide identity over at least 60% of the genome) were excluded (n=50), resulting in a total of 833 non-redundant MAGs representing characterized mouse gut microbes.

150 out of the 853 MAGs were present in the pre-FMT mice with a mean relative abundance greater than 0.1% (Figure 1a), consisting of predominantly Firmicutes bacteria (Figure 1a). Collectively, these MAGs explained a substantially larger fraction of mWGS reads than PanDB, a microbial reference database consisting of the pangenomes of approximately 20,000 microbial species deposited in GenBank ^23^ (Figure 1b), representing a significant improvement in mouse microbiome profiling compared with previous studies. Based on this improved compositional profile, we found that the microbial compositions of the pre-FMT huCD34 and SGM3 mice were significantly different (Figure 1a and 1c, PERMANOVA based on Bray-Curtis distance p=0.02), although the alpha diversities within each community were comparable for huCD34 and SGM3 mice (Figure 1c), suggesting that the humanized huCD34 mice and the immunodeficient SGM3 mice host distinct microbial communities with a comparable diversity.

**Figure 1.**
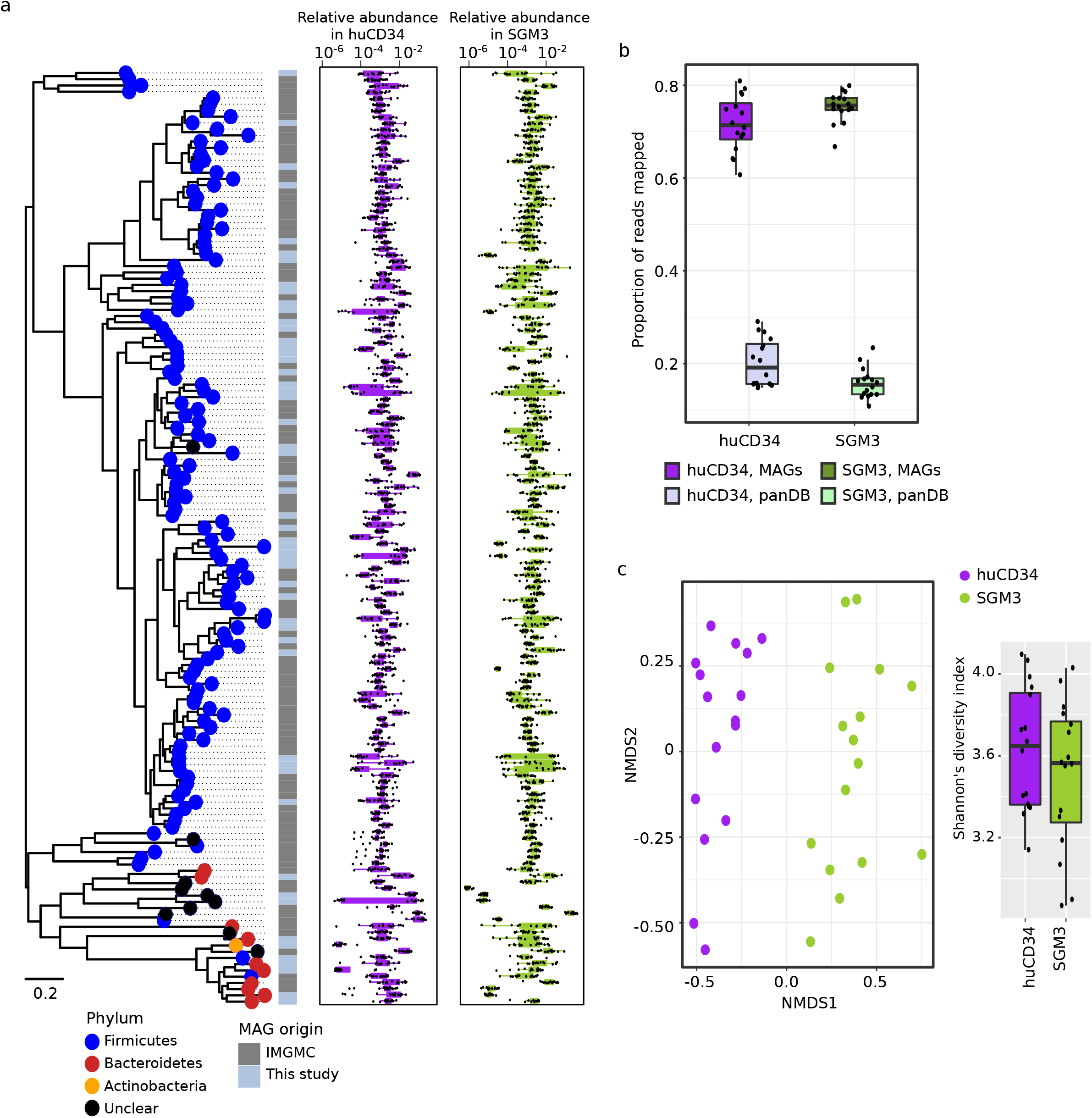
Pre-FMT mouse microbiome reconstructed using reference and *de novo* MAGs. a, Phylogenetic relationship and relative abundances of 150 high quality MAGs that constituted the pre-FMT microbiome of the huCD34 and SGM3 mice. b, the proportion of mWGS reads mapped to the panDB database and the high-quality MAGs. c, diversity of the pre-FMT mouse microbiome based on the relative abundances of the MAGs. left, multidimensional scaling of the pre-FMT microbiome samples based on the relative abundance profiles; right: Shannon’s diversity index based on the relative abundance profiles.

We do note that the huCD34 and SGM3 mice were grown in different facility rooms before being sent to the same room for FMT experiments with a week acclimation; thus the observed differences could have resulted from historic room effects in addition to the effects of humanization. Therefore, in this study we focused on microbial colonization dynamics after antibiotic clearance and FMT, but not the pre-FMT microbiome compositions.

### Transplantation of diverse healthy human fecal microbiota (FMT) into the mice

To compare the effect of mouse features, including immunocompetence and humanization, on the colonization dynamics of exogenous microbes during FMT, we combined the present data with our previous datasets addressing FMT dynamics in C57BL/6J, an immunocompetent strain, and NSG mice (Figure 2a, Supplementary Table 2). These mice were treated with the same FMT and sampling procedures in the Jackson Laboratory. The same FMT donor sample was used for all experiments, and were sequenced whenever a FMT experiment was carried out (Figure 2b). The comparison of FMT dynamics in huCD34 with SGM3 and C57BL6 mice was of special interest, because SGM3 and C57BL/6J mice each differs from huCD34 mice in a different aspect (SGM3 mice have the same genetic background with huCD34 but are immunodeficient and C57BL6 mice are immunocompetent but have no human components in the immune system).

**Figure 2.**
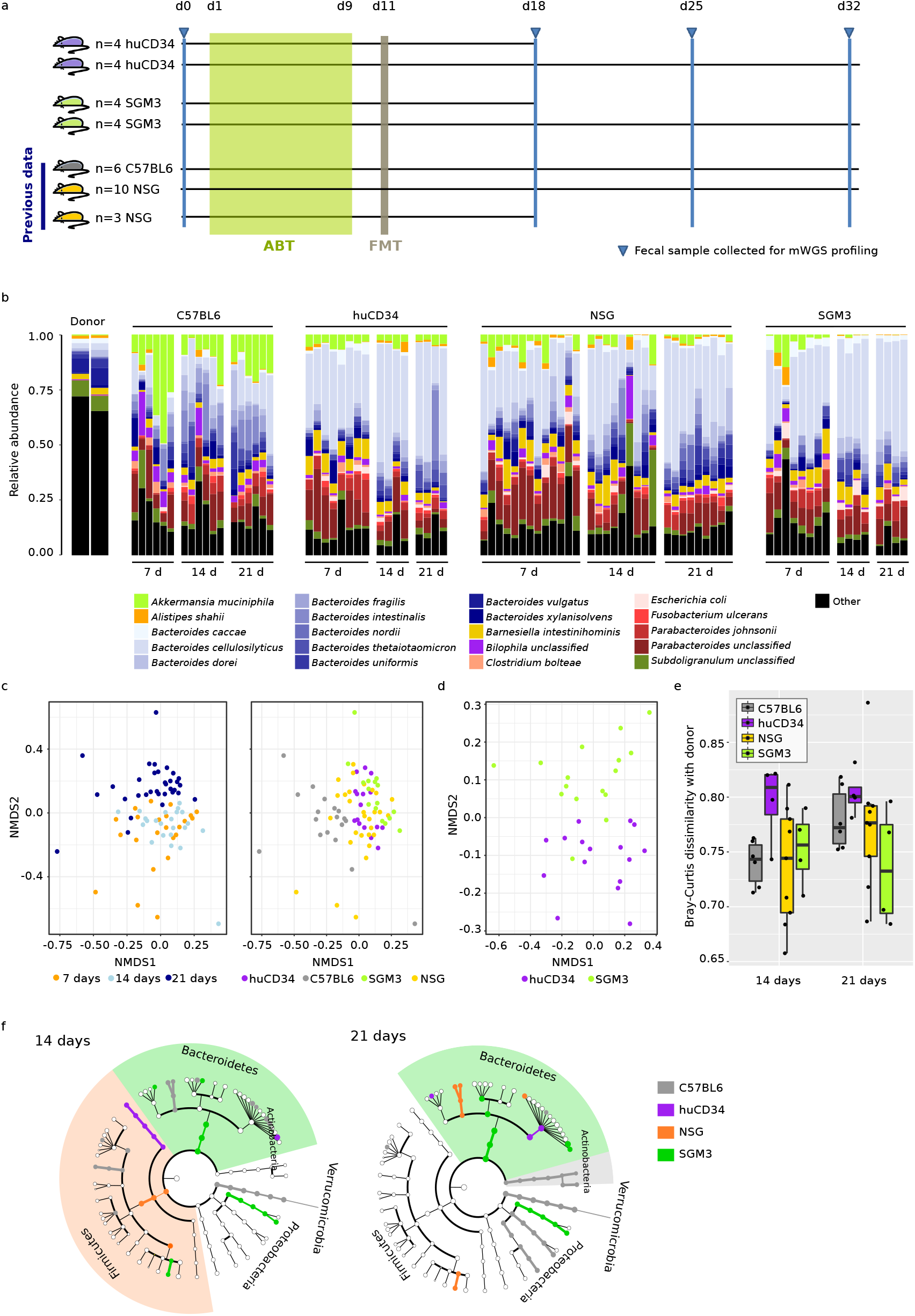
Taxonomic profiling of the post-FMT gut microbiome. **a**, the timeline of treatments and data collection. b, the proportion of mWGS reads mapped to the 20 most abundant microbial species in the post-FMT microbiome samples. The two donor samples were sequencing replicates of the same high-complexity donor community combined from six individual human fecal samples: the right replicate was sequenced during the FMT experiment carried out for the huCD34 and SGM3 mice, while the left replicate was sequenced during the FMT experiment carried out for the C57BL6/J and NSG mice. c, multidimensional scaling based on pairwise Bray-Curtis dissimilarities of the post-FMT microbiome samples. Samples are grouped and color-coded based on mouse type or sampling time (days post-FMT). d, multidimensional scaling based on pairwise Bray-Curtis dissimilarities of the post-FMT huCD34 and SGM3 microbiome samples. e, mean Bray-Curtis dissimilarity of the post-FMT microbiome samples to the donor samples. f, human fecal microbes enriched in different types of mice after FMT.

The transplantation of human microbiota into antibiotic-treated (ABT) mice was largely efficient regardless of the mouse type. To show this we reconstructed the composition of all post-FMT metagenomic samples across the four types of mice (C57BL6, NSG, SGM3 and huCD34). 65.0-66.8% of the donor microbial species (in relative abundances) can successfully colonize the mouse gut (i.e., with an average relative abundance > 0.1% among post-FMT mouse samples). The most successful colonizers (i.e. the 20 most abundant species in the post-FMT samples) consisted predominantly of bacteria from the Bacteroidetes phylum (Figure 2b). Similar to previous observations, recipient communities changed significantly over time (p<0.001, PERMANOVA on Bray-Curtis distance), with the community observed seven days post-FMT significantly different from the two later time points (Figure 2c, p<0.001 for 7 days vs 14 days/21 days, p=0.34 for 14 days vs 21 days, Benjamini Hochberg adjusted pairwise PERMANOVA on Bray-Curtis distance), suggesting that the colonized community could take at least a week to stabilize.

In addition, the recipient communities in the four types of mice differed after adjusting for the time effect (p<0.001, PERMANOVA on Bray-Curtis distance), suggesting an ecological selection specific to mouse strain that shaped the post-FMT gut community. C57BL/6J mice had the most distinct post-FMT microbiome (Figure 2c, p<0.001, Benjamini Hochberg adjusted pairwise PERMANOVA on Bray-Curtis distance), characterized by Verrucomicrobia, including *Akkermansia muciniphila*, which were consistently associated with the C57BL/6J mice (Figure 2f, linear discriminant analysis p<0.05). Also, humanization by HSC transplantation did appear to impact microbial colonization: the post-FMT microbiome in huCD34 significantly differed from the SGM3 mice (Figure 2d, p=0.03, Benjamini Hochberg adjusted pairwise PERMANOVA on Bray-Curtis distance), characterized by Bacteroidetes bacteria consistently associated with SGM3 mice (Figure 2f, linear discriminant analysis p<0.05). Interestingly, although huCD34 mice has, presumably, an immune system more similar to human, the post-FMT community in huCD34 was not more similar to the donor fecal microbiome (Figure 2e) – in fact trending towards more divergent from the donor community compared to the other three types of mice, though lacking in consistent microbial markers to differentiate them (Figure 2f). Collectively, these results suggested that immune selection may not be the predominant determinant of the fate of the colonizing microbes, although an effect of the engrafted human immune cells (SGM3 vs. huCD34) or, more generally, human-specific immune features (SGM3 vs. C57BL/6J and NSG) was observed.

### Dynamics of engrafted huCD34 immune components are nonspecific to human FMT

Having observed an effect of the engrafted human immune components on the post-FMT microbial community, we then investigated the converse, whether the engraftment of human microbiota affected composition and frequency of major human immune cell types one and three weeks after FMT (Figure 3a). We examined counts and frequencies in the bone marrow and spleen as primary reservoirs for immune maturation and performed transcriptomics on mouse peripheral blood leukocytes. Surprisingly, the introduction of human microbiota did not result in differences in the frequencies or numbers of human immune cell types. In both the spleen and bone marrow, FMT had no significant impact (p=0.229, ANOVA) on the frequencies of human leukocytes (i.e. the proportion of huCD45+ among all CD45+ cells, Figure 3b). Within the human leukocytes, the composition of the human leukocytes showed no significant differences between FMT and noFMT mice (Figure 3b, p=0.127 for spleen and p=0.399 for bone marrow, PERMANOVA based on Bray-Curtis distance). These results suggested that the engrafted human leukocytes in huCD34 are unresponsive to human-specific microbiota, at least within the first 21 days post-FMT.

**Figure 3.**
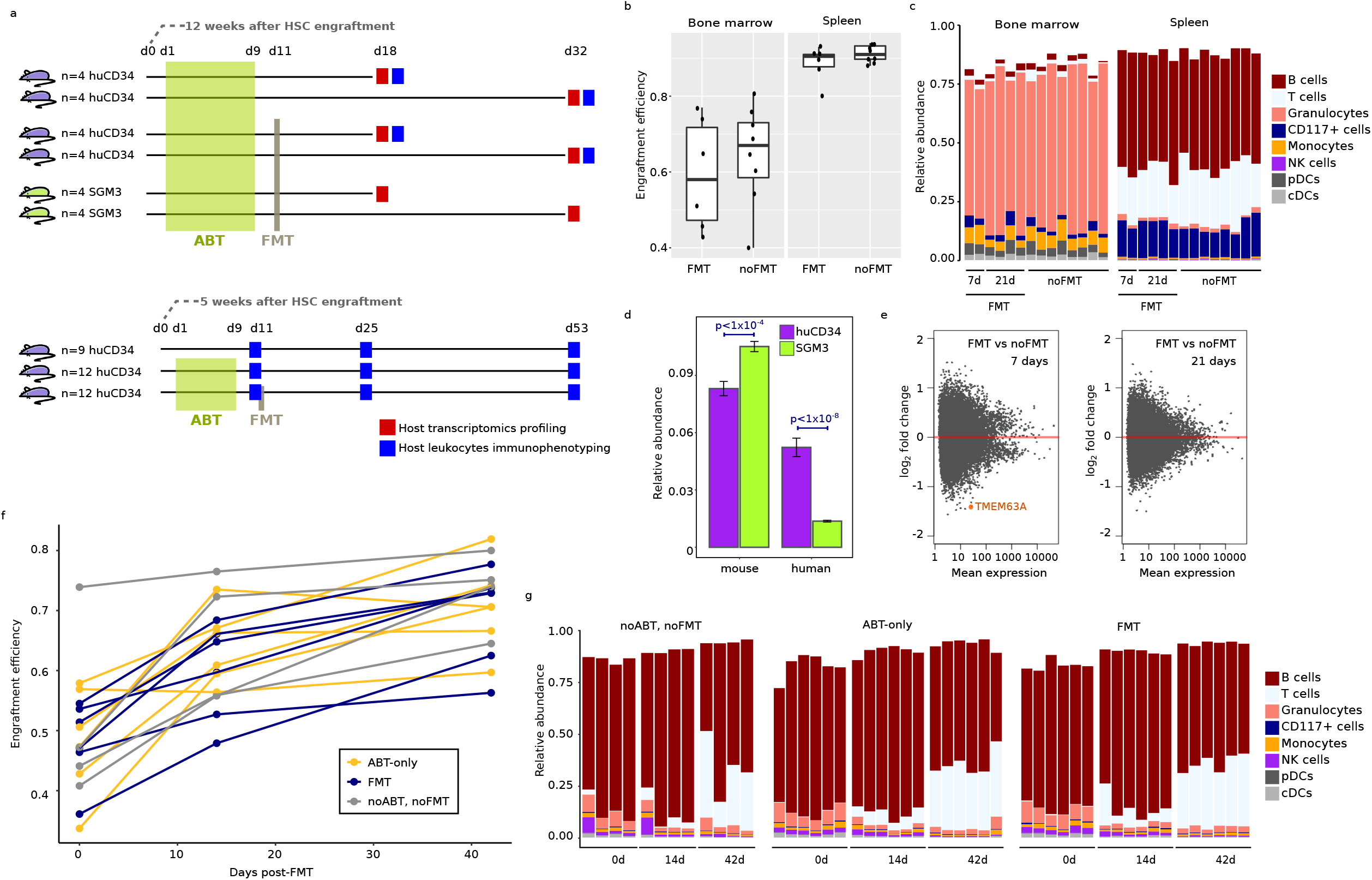
Engrafted human leukocytes are resilient to microbiome perturbation. **a**, the timeline of treatments and data collection. b, the engraftment efficiency of the human CD45+ cells in huCD34 mouse bone marrow and spleen, with or without FMT. Due to the lack of response, data from seven and 21 days post-FMT were combined to form the “FMT” group. c, relative abundances of different types of human leukocytes in the huCD34 mice. d, the proportion of Quantseq reads that mapped to the human reference genome (GRCh28) or the mouse reference genome (mm10). e, enrichment/depletion of human transcripts in the huCD34 mice post-FMT. TMEM63A, the only transcript showing a statistically significant difference, was highlighted. f, the engraftment efficiency of human leukocytes in the peripheral blood of the huCD34 mouse. ABT and/or FMT was conducted 6 weeks after the engraftment of human CD34+ cells. g, relative abundances of different types of human leukocytes in the huCD34 mice.

These observations were consistent with the transcriptomic profiles of the mouse peripheral blood. The huCD34 mice exhibited a significantly greater fraction of human transcripts compared to the SGM3 mice (Figure 3d, p<10^−4^, Wilcoxon rank-sum test), validating the successful engraftment of human leukocytes in huCD34 mice. Nonetheless only one transcript mapping to the TMEM63A gene showed significantly different levels (adjusted p<0.1, DESeq2) between the FMT and noFMT huCD34 mice (Figure 3e), potentially due to the modest sample sizes. Moreover, this difference was only significant 7 but not 21 days post-FMT (Figure 3e). Indeed, the TMEM63A gene was most prevalent in leukocytes including CD4+ and CD8+ cells, and was highly expressed in the gastrointestinal tract ^24,25^, consistent with its potential response to FMT. However, the lack of significant transient or long-term transcriptional changes between FMT and noFMT huCD34 mice suggested that the engrafted human leukocytes was functionally stable irrespective of whether a complement of largely human or largely mouse microbiota repopulate the gut after antibiotic treatment, at least within the first 21 days. The lack of a response of the engrafted human leukocytes to human FMT could be due to 1) FMT can modulate the process of hematopoiesis, but not a stable human leukocyte compartment established after hematopoiesis, 2) the effect of ABT has confounded the effect of FMT, and 3) the effect of FMT required more than 21 days to appear statistically significant. Therefore, we then monitored the long-term immunological influence of FMT, conducted during an early pulse of hematopoiesis, and in the meantime characterizing the potential influence of ABT (Figure 3a). To increase sensitivity, we tracked the immune cell compositions in the same huCD34 mice over time via RO bleeding at 0, 14, and 42 days post-FMT. The engraftment efficiency showed a significant increase over the post-FMT period (Figure 3f, p<0.0001, ANOVA after corrected for autocorrelation), suggesting that our experiment can capture the immunogenic effect of microbiome perturbation, or the lack thereof, during the process of hematopoiesis. However, consistent with the short-term experiments, neither ABT nor FMT exhibited any statistically significant influence on the engraftment efficiency (Figure 3f, p=0.46, ANOVA after corrected for autocorrelation). In addition to overall engraftment efficiency, FMT or ABT did not appear to alter the frequencies of human leukocyte lineages (Figure 3g, p=0.72, PERMANOVA based on Bray-Curtis distance), although these frequencies varied significantly over time (Figure 3g, p<0.001, PERMANOVA based on Bray-Curtis distance). These results again demonstrated the stable engraftment of human leukocytes despite substantial alterations in the gut microbiome, suggesting that huCD34 mice may not be sensitive enough to model a host-specific, microbiome-immune interaction.

### FMT dynamics and mouse-type dependency of Bacteroides strains

Previous studies have shown that conspecific bacterial strains had distinct colonization abilities depending on the host gut environment ^11,26^, consequently introducing diverse functional impacts on the host - an emerging topic in both microbial ecology and clinical microbiology. Given the importance of strain diversity, we resolved our mWGS data at strain resolution to compare strain-level colonization patterns across time points and mouse types. Specifically, we asked 1) if the post-FMT strain populations reflected the strain compositions in the donor population, 2) if different strains can co-exist in a post-FMT population, and 3) if the strain composition in a given post-FMT population could be time- or mouse-type-specific.

We first explored the strain-level colonization dynamics of four human fecal Bacteroides species (*B. xylanisolvens, B. dorei, B. ovatus, and B. vulgatus*), which were successful colonizers of the mouse gut, have important roles in the host ranging from immunomodulation to opportunistic pathogenicity ^27,28^, and had adequate sequencing coverage (mean per base read coverage of at least 2) in at least half of the post-FMT samples (n>42) for *de novo* strain inference. Predicting strain haplotypes based on single nucleotide polymorphisms and using frequencies to infer strain relative abundance ^26^, we found that the strain composition of each Bacteroides species differed across samples (Figure 4a). Surprisingly, terminal strain compositions were substantially different from the donor populations – the dominant strains (i.e. strains with the highest relative abundances) in the donor samples were rarely the dominant strains colonizing the mouse gut 21 days post-FMT (Figure 4a). This is in contrast to a recent human-human FMT study ^26^, in which donor strain abundance was shown to be the major determinant of colonization success. This discrepancy underscores the remarkable difference between the human and mouse gut environments, humanized or not, and suggests that even a highly humanized mouse model such as the HMA huCD34 might not faithfully reflect the microbial interactions in a human host.

**Figure 4.**
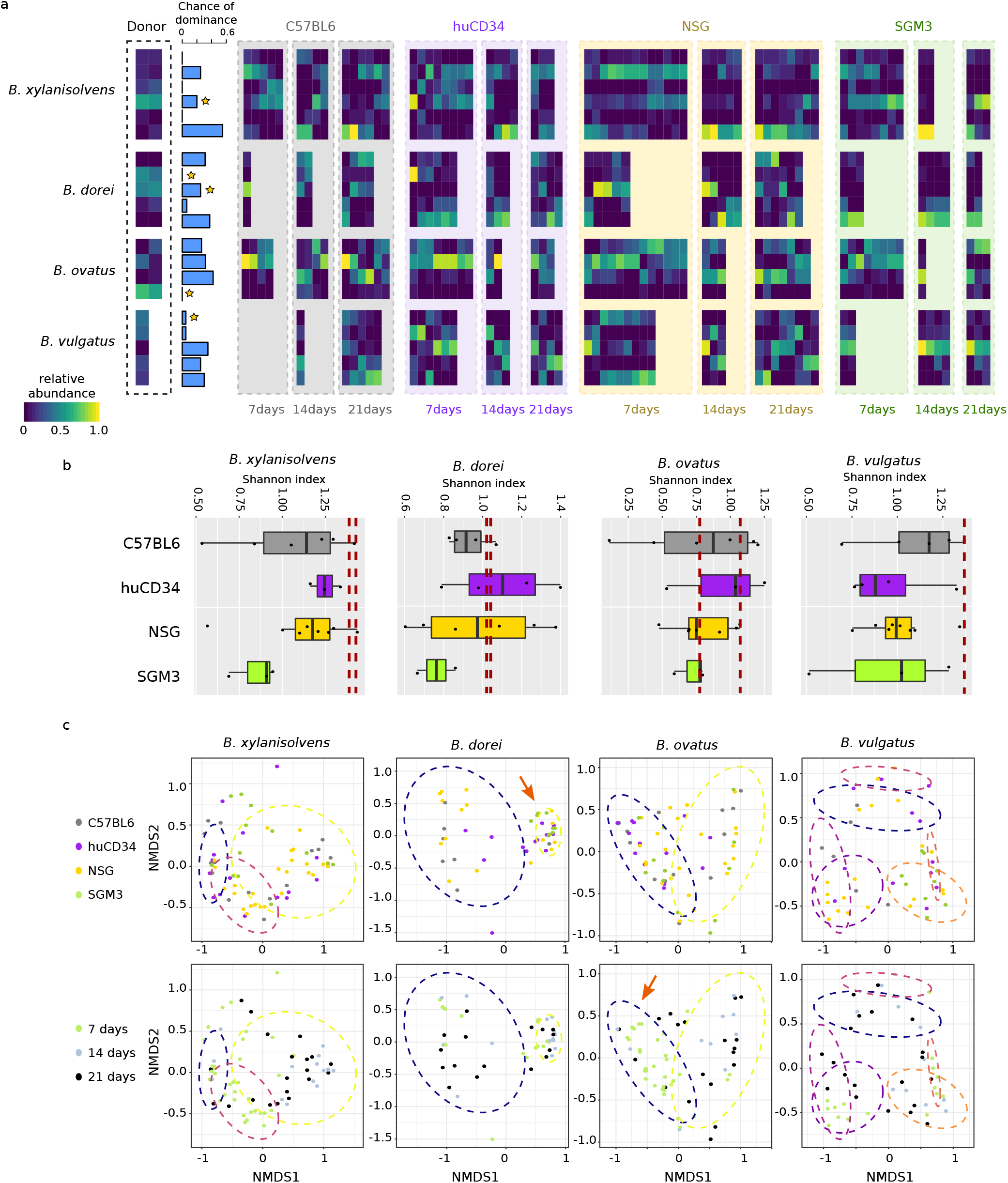
Strain-level colonization dynamics of human fecal Bacteroides species in the mouse gut. **a**, the within-species relative abundances of the conspecific strains identified using StrainFinder. A missing column indicated that a sample did not pass the filtering step of StrainFinder for the species of interest. “Chance of dominance” of a strain was estimated based on the proportion of post-FMT samples where the strain had higher relative abundance than any other conspecific strain. Strains that were dominant in at least one donor sample (a sequencing replicate of the same high-complexity donor community) were marked with a star. b, the strain-level diversity (Shannon’s diversity index) of Bacteroides species 21 days post-FMT (box plots), compared to the strain-level diversity in the donor samples (dashed lines). c, association between population states (strain relative abundance profiles) and mouse type or sampling time. Population states identified using Gaussian-mixture-model clustering were indicated with ellipses. Significant associations were indicated with red arrows.

We also found that different conspecific strains can co-exist in the post-FMT population (Figure 4a). While we cannot determine how many strains were derived from each individual contributing to the FMT donor sample (representing original strain interactions), one potential reason for strain coexistence could be the lack of strong purifying selection in the mouse gut, which would otherwise result in a decrease in strain-level diversity. Instead, we observed that the strain-level diversities in the post-FMT communities that were often as high as the donor sample (Figure 4b, Benjamini-Hochberg corrected Wilcoxon rank sum test on Shannon’s diversity index between the donor and the post-FMT populations, p≥0.2 for any Bacteroides species in any given mouse type). However, taking into account statistical power, purifying selection could be present for *B. xylanisolvens* strains and *B. vulgatus* strains; their post-FMT diversities were consistently lower than the donor sample in all mouse types (Figure 4b, p=0.05 for *B. xylanisolvens* and p=0.09 for *B. vulgatus*, Wilcoxon rank sum test with all mouse types combined). These results demonstrated that 1) unlike in human studies, Bacteroides populations were reshaped during FMT in the mice, causing substantial divergence from the human donor samples, and 2) although abundances of Bacteroides strains were reshaped by the mouse gut environment, different conspecific strains were still allowed to coexist in the novel environment.

We then investigated potential determinants of this post-FMT strain-level reshaping. Strain composition in the post-FMT populations were dependent on time (p<0.05 for all species except *B. dorei*, PERMANOVA based on Bray-Curtis distance) and mouse type (p<0.01 for all species), showing both the dynamic nature of strain composition and their potential response to different host features. To describe such alternative strain compositions, we clustered strain composition profiles for each Bacteroides species using a gaussian mixture model with model selection based on the Bayesian information criterion (BIC) (similar to Sirota et al., 2013). We assumed that, for a given species, each cluster of strain compositions represented a distinct population state; associations between population states and time point or mouse type might indicate a strain-level response to temporal or mouse-specific features. We found that *B. dorei* populations exhibited distinct population states when colonized in SGM3 mice (Figure 4c, p=0.017, estimated by unrestricted permutation). Although the underlying mechanism is unclear, it is surprising that strain compositions of *B. dorei* in SGM3 mice were more consistent across samples compared to NSG mice, while the two types of mice were highly similar genetically, suggesting a potential interaction with the transgenic human components in the SGM3 mice.

In terms of temporal patterns, we observed strain populations that could be highly dynamic or transient. *B. ovatus* populations were highly dynamic, shifting from a starting state observed at seven days post-FMT to an alternative state observed at later time points (Figure 4c, p<0.001, estimated by unrestricted permutation). Interestingly, the trajectory of the shifts appeared to be consistent in all four mouse types, demonstrating an example of strain-level dynamics robust to host features. Consistent with the temporal changes observed at the species level, the observed strain-level dynamics showed that human fecal strains could take at least a week to stabilize in the mouse gut.

### FMT dynamics and mouse-type dependency of Akkermansia muciniphila strains

Next, we interrogated the strain-level dynamics and mouse type specificity of *A. muciniphila*, a microbe known for its inverse association with obesity, diabetes, inflammation, and resistance to cancer immunotherapy ^30–32^. *A. muciniphila* was abundant in only a subset of mice pre-FMT (all huCD34 and C57BL6 mice, about half of the NSG mice, and no SGM3 mice), and human-associated *A. muciniphila* only colonized well in a subset of mice (all huCD34 and C57BL6 mice and about half of the NSG mice). We found that the pre-FMT mouse-associated *A. muciniphila* abundance was an important determinant of the colonization success of human-associated *A. muciniphila* post-FMT (adjusted R^2^=0.50 and slope different from zero p=9.8×10^−5^, log-log regression with all mouse types combined, Figure 5a). This suggests that pre-FMT *A. muciniphila* may have shaped the mouse gut environment rendering it permissive to colonization of other strains, for example by altering nutrient availability. To further investigate the population structures of *A. muciniphila*, we inferred the pre- and post-FMT strain compositions of *A. muciniphila* using a reference-based method, in which sequencing reads were mapped and assigned to the closest reference genome in NCBI GenBank (Supplementary Table 3). We did not infer strain compositions *de novo* because unlike the Bacteroides species, *A. muciniphila* did not have sufficient read coverage for *de novo* haplotype inference. We found that pre-FMT *A. muciniphila* strain compositions were strikingly diverse across mouse types (Figure 5b). Notably, all mouse samples were then dominated by a single *A. muciniphila* strain type from the FMT donor (Figure 5b), which represented a close relative of the GenBank strain GCF_002885555. Surprisingly, the dominant strain in the post-FMT samples was not the dominant strain in the donor fecal samples (Figure 5b), demonstrating a strong purifying selection imposed by the mouse gut environment. We hypothesized that the combination of modified environment by the mouse *A. muciniphila* and the unique functional qualities of the human *A. muciniphila* favored the dominance of this single strain type. We thus investigated potential functions that differed between the pre-FMT and post-FMT *A. muciniphila* populations with the hypothesis that mouse-associated *A. muciniphila* may possess unique functions to facilitate primary colonization, and the human *A. muciniphila* possess unique functions that allow its dominance. We assembled and annotated *A. muciniphila* genes *de novo* from the mWGS data and compared their abundances between pre- and post-FMT samples. Interestingly, pre-FMT *A. muciniphila* in most huCD34 and NSG mice had different gene contents compared to C57BL/6J mice (Figure 5c). The difference could be attributed to two groups of genes, group I and group V (Figure 5c). While most of the genes in these two groups could not be annotated, group I genes, enriched in C57BL6 mice, contained a large fraction of ion transport and metabolism genes (Figure 5c), potentially related with the immune control of intestinal ion transportation ^33^. Strikingly, pre- and post-FMT *A. muciniphila* exhibited a significantly different gene set, underscoring the difference between *A. muciniphila* strains of mouse and human origins. Post-FMT *A. muciniphila* had relatively fewer accessory genes compared to pre-FMT *A. muciniphila*, suggesting that the post-FMT *A. muciniphila* population also had lower strain-level genetic diversity. Despite the low genetic diversity, a cluster of genes (group III) were abundant only in the post-FMT *A. muciniphila*. Given that the post-FMT *A. muciniphila* can colonize both human (FMT donor) and mice (FMT recepients), group III genes could be associated with diversification of colonization abilities in diverse host species. This group of genes were mostly metabolism-related and showed an enrichment of genes involved in carbohydrate metabolism and transport (p=0.066, Benjamini-Hochberg adjusted). This suggests that the colonization ability of *A. muciniphila* in different host species may be motivated by different aspects of metabolic versatility. Overall, our results demonstrated that, while strain co-existence was common for the Bacteroides species, the post-FMT *A. muciniphila* population was swept by purifying selection, resulting in consistent dominance of a single strain type.

**Figure 5.**
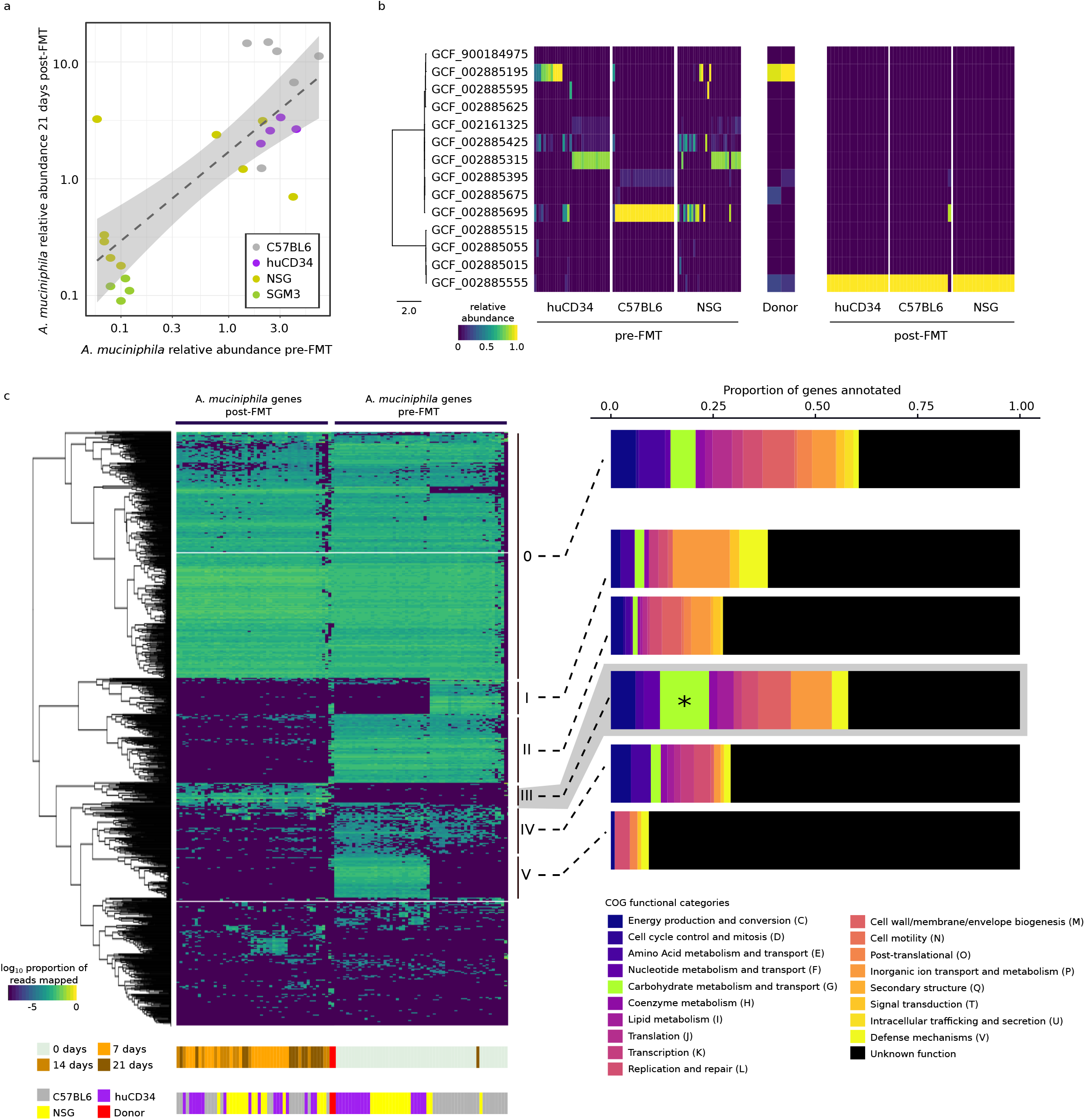
Strain-level dynamics of *A. muciniphila* in the mouse gut. a, the association between the pre-FMT and the post-FMT (21 days) relative abundances of *A. muciniphila*. b, the within-species relative abundances of the *A. muciniphila* strains in the pre- and post-FMT samples. Samples from SGM3 mice were excluded for the consistently low *A. muciniphila* abundances in these mice (Figure 5a). c, the gene content diversity in *A. muciniphila* strains pre- and post-FMT. Six groups of genes with distinct abundance distributions were annotated. The COG functional category (G) that was significantly enriched in group III was marked with a “*”.

## Discussion

One of the central roles of the microbiome is its interactions with the immune system. Humanized mouse models such as HMA mouse models and huCD34 mouse models have been developed to respectively simulate the human microbiome and human immune functions. We hypothesized that simultaneous humanization of both components might be suitable for studying human immune-microbiome interactions. In this prospective study, we combined the two humanization procedures to generate the HMA huCD34 mouse model – a mouse model with engrafted human leukocytes and human microbiome introduced through FMT. We found that the post-FMT microbiome in huCD34 mice was significantly different from C57BL/6J, NSG, and SGM3 mice, suggesting distinct immune selection imposed by the engrafted human leukocytes. Interestingly, the human FMT microbiome did not appear to have a reciprocal impact on the engrafted human leukocytes, both in terms of abundances and compositions of cell types or gene expression profiles. In addition, such immune selection by the humanized immune system did not render the transplanted microbiome more similar to the donor microbiome, suggesting that the donor microbiome could be shaped by factors additional to immune selection.

By resolving the post-FMT microbiome at strain-resolution, we identified different factors that could shape engraftment and dynamics of the donor microbiome. We tracked the colonization dynamics of bacterial strains in the different types of mouse models. In a surprising divergence from results from human-to-human FMT experiments ^26^, colonization success of strains in the mice, including the humanized mice, were not determined by their abundances in the original FMT donor. For example, while donor Bacteroides strains often co-existed in the post-FMT microbiome in the mice, their compositions significantly diverged in the post-FMT microbiome, showing mouse-type- or time-point-dependent population structures. On the contrary, a single type of donor *A. muciniphila* strain consistently dominated all post-FMT samples regardless of the time of sampling, demonstrating strong purifying selection common to all mouse types.

A major conclusion from this study is that although a variety of humanized mice have been developed to model different facets of human biology (e.g., the human immune system or the human microbiome), they might not faithfully reproduce the interactions between these factors. This is illustrated by our findings that the human immune system modeled in the huCD34 mice was largely unresponsive to the human-associated microbes introduced via FMT, irrespective of stage of immune maturation (Figure 3), and the post-FMT microbiome in huCD34 mice was not more similar to the donor microbiome compared to the non-humanized mice (Figure 2e). The underlying reasons for this observed limitation could be one of the following four factors. First, humanized mice are imprecise for the aspects of human biology they were designed to model. For example, the engrafted immune cells in huCD34 mice cannot be an exact simulation of the human immune system – they have impaired IgG production, immature NK cells, and a contamination of mouse granulocytes, macrophages, and DCs in the immune cell populations, in addition to others ^20^. Also, NSG and SGM3 mice have a targeted mutation in the IL-2 receptor common gamma chain, which significantly enhances the engraftment of human immune cells but also can result in a lower level of immune cell migration to the intestine, which could limit interactions with the human microbiota ^34^. Similarly, we found that the post-FMT HMA mice typically host only half of the donor fecal microbes, whether due to technical reasons such as the loss of fecal microbes during sample processing, or biological reasons such as the importance of the order of ecological succession and selective forces in the mouse gut.

Second, human-specific immune-microbiome interactions might involve other factors of human biology that are not modeled in the humanized mice. Factors such as diet, lifestyle, metabolic rate, hormones, and the nervous system are known to impact both the immune system and the microbiome ^35–38^ and therefore could play a role in regulating the immune-microbiome interaction. In addition, donor-to-donor variability in the cord blood is a common confounder in generation of the huCD34 models ^20^. These factors can be incompletely accounted for in present mouse models. Third, due to the engraftment into a foreign environment and exposure to foreign microbes, the human leukocytes could be unresponsive due to mechanisms such as immune exhaustion ^39^ – a phenomenon often observed when T cells are persistently exposed to antigens or inflammatory signals.

Finally, true immunological differences could be obscured by low statistical power. The major limiting factor in statistical power was the sample size of the huCD34 mice, because the production of huCD34 mice is costly and slow. For example, in the short-term experiment (Figure 3a), huCD34 mice were generated following a standard procedure: after CD34+ engraftment, the mouse cohort is then raised for 12 weeks before testing for human leukocyte engraftment efficacy, and individuals with <0.25 engraftment efficiency (which can represent a significant fraction of the original cohort) are excluded from downstream experiments. Even for a longer-term experiment without cohort selection, each experiment was controlled for cord blood donor, a known variate, and a donor can only be used to create a limited number of mice. This sample size bottleneck should also be taken into consideration when designing any microbiome-related experiments using huCD34 mice (whether or not HMA), as comparative microbiome analyses often require large sample sizes to achieve sufficient power, even for biologically meaningful effect sizes.

Given the differences between humans and mice, human-to-mouse FMT is a process of “ecological fitting”, where exogenous colonizers attempt to invade, then persist in a novel environment with their pre-existing suite of traits ^40^. Differing from human-to-human FMT studies where human microbes re-colonize an environment to which they are already adapted, human-mouse FMT exhibited distinct colonization dynamics, even at strain resolution. By tracking the dynamics of the Bacteroides strains, we found that the post-FMT strain abundances did not correlate with the donor strain abundances, suggesting strong environmental pressures that shaped the colonizing strain populations in the mice. This finding contrasted a human-human FMT study, where strains were resolved using the same computational methods: strain abundances in the donor samples appeared to be the major determinant of colonization success ^26^. This difference between FMT outcomes underscored the impact of heterologous host selection on strain colonization dynamics. Note that the host environmental differences in the two studies also include the presence/absence of a native gut microbiome: in the human-human FMT study, the donor strains need to compete with the native host microbiome. In this study, the native host microbiome was removed by ABT, which theoretically should have facilitated species/strain invasion. Indeed, we previously found that the absence of a native gut microbiome represented an additional source of selection, interspecies, and inter-strain interactions that can significantly influence the colonization outcome ^11^. Given the observed differences in strain colonization dynamics and potential underlying reasons, a following question will be whether these differences were due to heterologous hosts, the presence/absence of a native microbiome, or, more likely, both? Deciphering the mechanistic determinants of strain colonization outcomes are significant for microbial ecology, microbial pathogenesis, and probiotics design, yet this question is challenging to approach because high-dose ABT cocktails ought not be used in humans simply to investigate the underlying microbial ecology. Therefore, inference of colonization determinants might require robust longitudinal models generated from large comparative studies of FMT in mice with native microbiomes, mice without native microbiomes, healthy patients, and patients with dysbiotic microbiomes.

A striking finding in our study was the diversity of selective forces that shaped microbial populations at the strain level – these could be diverse and depended on both microbial and host features. For example, the observed coexistence of Bacteroides strains with a stable strain-level diversity, together with the consistent dominance of a single strain in the *A. muciniphila* population suggested that purifying selection was a driving force shaping the *A. muciniphila* population but not that of the Bacteroides species. In addition, the speed of microevolution can differ among microbial species. For example, strong purifying selection swept the *A. muciniphila* population within 7 days post-FMT, while *B. ovatus* showed a dynamic shift in population state from 7 days to 14 days post-FMT. While these strain-level dynamics were mouse-type-independent, colonization dynamics of *B. dorei* exhibited highly consistent strain compositions only when colonizing SGM3 mice. This mouse type-dependence indicated that the transgenic cytokines present in SGM3 but not NSG mice (which is otherwise genetically identical) could influence bacterial strain colonization. An association between gut microbes and cytokines has been identified in numerous health and disease states ^41,42^. However, these associations were predominantly identified in immunocompetent hosts, while our findings suggested that the interaction between the transgenic cytokines with the Bacteroides strains might be specific to an immunodeficient background, because the association was not observed in huCD34 mice that also carried the transgenic cytokines. This finding more generally underscores the complexity of the microbiome-immune interaction and the importance of probing the microbiome at strain level – a bacterial population state could respond differently to the presence of different subsets of immune components, and taken together with the different environmental characteristics of a host’s gut microenvironment, combine to present unique driving forces that shape population states at the finest taxonomic levels.

## Methods

### Overview of the study

The overarching goal of this study was to compare the effect of mouse features, including immunocompetence and humanization, on the colonization dynamics of exogenous microbes during FMT, and correspondingly, to understand if FMT affects the engrafted human immune cells. The experimental timeline is summarized in Figure 2a and Figure 3a.

### Construction of the HMA mice models

FMT was conducted as described previously ^11^. The experimental design was illustrated in Figure 2a (collection of microbiome data) and Figure 3a (collection of host transcriptomics and immunophenotyping data). Briefly, SGM3 (NOD.*Cg-PrkdcscidIl2rg*_*tm1Wjl*_Tg(CMV-IL3,CSF2,KITLG)1Eav/MloySzJ) and huCD34 mice (female SGM3 mice engrafted with human hematopoietic stem cells, or CD34+ cells, from a single cord blood donor at 5-6 weeks of age) were purchased from the Jackson Laboratory (Bar Harbor, ME, USA) and produced in JAX WEST (Sacramento, CA, USA). For ABT, a broad spectrum antibiotic cocktail (1mg/mL ampicillin; 5mg/mL streptomycin; 1mg/mL colistin; 0.25mg/mL vancomycin) was added directly into the drinking water and the drinking water was changed with fresh antibiotics once per week for two weeks. Healthy human stool samples were purchased from OpenBiome (Somerville, MA, USA) stool collection. For FMT, the mice were oral gavaged with 0.2mL/10g of stool sample (mixed from 6 healthy human donors) resuspended in glycerol. Note that, we used a mixture of human fecal samples to explore competitions between the microbial strains in different human donors. Therefore, the mixed samples were more diverse and did not mimic any single fecal sample.

### Effect of FMT on the engrafted human leukocytes in the huCD34 mice

Two sets of experiments were conducted to test the short-term and long-term effect of FMT on the engraftment of human immune cells (Figure 3a). For the short-term experiments, we only used huCD34 mice that had an engraftment efficiency of no less than 0.25 in the peripheral blood 12 weeks after the engraftment so as to measure the effect of microbiome perturbation on a relatively mature human immune system. Engraftment efficiency was calculated as the proportion of huCD45+ cells among all CD45+ cells. For the long-term experiments, we used all huCD34 mice 6 weeks after the engraftment regardless of the engraftment efficiency. This is to reveal the effect of microbiome perturbation *during* hematopoiesis, instead of *after* the maturation of human leukocyte lineages.

In the short-term experiments, half of the huCD34 mice (n=8) were sacrificed seven days post-FMT, while the other half (n=8) were sacrificed 21 days post-FMT, before tissue samples were collected from the spleen and bone marrow for immunophenotyping (described below), and blood samples (via Retro-orbital bleeding) were collected for transcriptomics profiling (described below). Half of these huCD34 mice (n=8) had received ABT but not FMT (Figure 3a) and were used as non-FMT controls. Additionally, eight ABT and FMT treated SGM3 mice were used as non-human controls to validate that the transcriptomics had captured the expression of human-specific transcripts in huCD34 (Figure 3a).

In the long-term experiments, 12 huCD34 mice were treated with both ABT and FMT using the same approach as described above, and their peripheral blood samples (via Retro-orbital bleeding) were collected right before FMT, 14 days post-FMT, and 42 days post-FMT. The peripheral blood samples were pooled in groups of 2-3 mice for immunophenotyping (Supplementary Table 4). As controls for FMT and ABT, 12 huCD34 mice were included that had received ABT but not FMT (ABT-only), and 9 huCD34 mice were included that had not received ABT or FMT (Figure 3a, Supplementary Table 4). All mice used in the long-term experiment were from the same cohort (female, 5-6 weeks of age before engraftment).

### Additional HMA mice metagenomics data

A shotgun metagenomics dataset on 6 C57BL6/J and 5 NSG mice described previously ^11^ (Supplementary Table 2, each mouse had stool samples collected at both 7 and 21 days post-FMT before sacrificed) was included in the study to compare the effect of mouse features on colonized microbes (Figure 2a). In addition, we included previously unpublished data on 8 NSG mice produced from both JAX WEST (Sacramento, CA, USA) and JAX EAST (Bar Harbor, ME, USA) (Supplementary Table 2) in which 4 were sacrificed 7 days post-FMT, and the other 4 were sacrificed 21 days post-FMT (Figure 2a). ABT and FMT with mixed human fecal samples were conducted using the same approach (as described above and previously in ^11^) for all mice. Stool samples from all mice were collected at the same time intervals, from which total metagenomic DNA was extracted and sequenced with the same procedure as described below (Figure 2a).

### Total DNA extraction from mouse stool samples

Collection of mouse stool samples and extraction of total metagenomic DNA were conducted as described previously ^11^. Mouse stool was collected into Cell & Tissue Lysis buffer (Ambion, Austin, TX, USA) and homogenized with a pestle before being frozen at -80°C. DNA was extracted using the Qiagen (Germantown, MD, USA) QIAamp 96 DNA QIAcube HT Kit with the following modifications: enzymatic digestion with 50 μg lysozyme (Sigma, St. Louis, MO, USA) and 5U each of lysostaphin and mutanolysin (Sigma) for 30 minutes at 37°C followed by beadbeating with 50μg 0.1mm zirconium beads for 6 minutes on the Tissuelyzer II (Qiagen) prior to loading onto the Qiacube HT. DNA concentration was measured using the Qubit high sensitivity dsDNA kit (Invitrogen, Carlsbad, CA, USA).

### Metagenomic shotgun sequencing

Library preparation and metagenomic sequencing were conducted as described previously ^11^. Illumina libraries were created using Nextera XT DNA Library Prep Kit (Illumina, San Diego, CA, USA) with reduced reaction volumes: 200pg of DNA were used (160pg/μL×1.25μL), and tagmentation and PCR reagent volumes were reduced to 1/4 of the standard volumes. Tagmentation and PCR reactions were carried out according to the manufacturer’s instructions. The reaction mixtures were then adjusted to 50μL by adding dH2O, and the AMPure (Beckman Coulter, Brea, CA, USA) Cleanup was carried out as per the manufacturer’s instructions. Libraries were then sequenced with 2×150bp paired end reads on an Illumina HiSeq2500. For quality control, expected vs. observed frequencies of species in sequencing of a bacterial mock community were closely matched (data not shown).

Sequencing adapters and low quality bases were removed from the sequencing reads using scythe (v0.994) ^43^ and sickle (v1.33) ^44^, respectively, with default parameters. Host reads were then filtered by mapping all sequencing reads to the hg19 human reference genome or mm10 mouse reference genome using bowtie2 (v2.2.8) ^45^, under “very-sensitive” mode. Unmapped reads were used for downstream analyses. Characteristics of the shotgun metagenomic sequencing data were summarized in Supplementary Table 2.

### De novo assembly and binning

To enhance taxonomic profiling of the mouse native microbiome samples, draft quality genome assemblies were inferred based on the mWGS data using a binning strategy, as previously described in ^11^. Sequencing reads from mouse samples before receiving ABT or FMT were pooled for *de novo* assembly using MEGAHIT (v1.0.6) ^46,47^ with default parameters. The resulting contigs were filtered by size (>1000bp) and sequencing reads were mapped back to the contigs using bowtie2 (v2.2.8) ^45^ under “very-sensitive” mode. Genome bins were constructed from the contigs using MetaBat ^48^ with the runMetaBat.sh wrapper using default parameters, accepting genome bins with mean depth of coverage >=1 in each library. The resulting genome bins were estimated for genome completeness and co<u>ntamination u</u>sing the CheckM (v1.0.9) “lineage workflow” pipeline with default parameters ^49^. Genome bins with at least 90% completeness and at most 5% contamination (i.e., high quality MAGs) were included for downstream analyses. These high quality MAGs were assigned taxonomic labels (Supplementary Table 1) using the CheckM (v1.0.9) “lineage workflow” pipeline ^49^. For additional taxonomic resolution, 830 high quality MAGs (i.e. the IMGMC MAGs) reported by Lesker et al^22^ were retrieved from github.com/tillrobin/iMGMC. Duplicated MAGs, defined as MAGs that belonged to the same species, were identified using the dRep pipeline (v2.0.0, the “cluster” module with default parameters) ^50^. Duplicated IMGMC MAGs with an average nucleotide identity of at least 96.5% ^51^ over at least 60% of the genome bases ^52^ to a MAG generated de *novo i*n this study were excluded. The resulting set of de-duplicated MAGs were combined into a reference catalog; mWGS reads of the pre-FMT samples were then mapped to this catalog using bowtie2 (v2.2.8) ^45^ under “very-sensitive” mode. The fraction of mWGS reads mapped to each MAG were computed using samtools (v1.8) ^53^. Extremely rare microbial species – species with an average relative abundance <0.1% – were excluded from the profiles prior to downstream analyses ^11^.

### Taxonomic composition profiling of post-FMT samples

Taxonomic compositions of the post-FMT mWGS samples, and the human donor samples, were profiled using MetaPhlAn2 (v2.5.0) ^54^ as described previously ^11^. Extremely rare microbial species – species with an average relative abundance <0.1% – were excluded from the profiles prior to downstream analyses. To illustrate the diversity between the microbiome samples, multidimensional scaling was conducted using the metaMDS function in the R package vegan ^55^. Human fecal microbes that were differentially enriched in different host conditions were identified using LEfSe ^56^(with the argument –o 1000000 as a normalization factor) based on the filtered and rescaled compositional profiles.

### Immunophenotyping

The procedure for preparing cell suspensions from different types of mouse tissues was described in Supplementary Method 1. The resulting cell suspensions were washed with 4 ml protein-free PBS, spun at 500xg for 5 min at 4°C before the supernatant was aspirated. The cells were resuspended in 1 ml protein-free PBS, 500µl of which were transferred to a 1.5ml Eppendorf tube, pelleted at 500xg with supernatant aspirated. 10µl Zombie UV fixable viability dye (BioLegend, San Diego, CA, USA) were added to the cells, before they were resuspended and incubated for 20 min at room temperature in the dark. The cells were then washed twice with 1ml FACS buffer (PBS + 2%FBS + 2% NaN_3_) and pelleted. 25µl of human panel antibodies A (Supplementary Table 5) were added to the tubes. The tubes were incubated for 30 minutes at room temperature in the dark, washed twice with 1ml FACS buffer and the cells were pelleted. After this 25µl of human panel antibodies B (Supplementary Table 5) were added to the tubes, before the tubes were incubated for 30 minutes at room temperature in the dark, washed twice with 1ml FACS buffer and the cells were pelleted. For fixation, 100µl BD fixation buffer (BDBiosciences, San Jose, CA, USA) were added to the resuspended cell pellets, followed by a 20-minute incubation at room temperature in the dark. The cells were washed twice with 1ml FACS buffer, pelleted and resuspended in 250µl FACS buffer. The cells were analyzed on a LSRII flow cytometer (BDBioscience, San Jose, CA, USA). An example illustrating the gating strategy used to identify different subsets of leukocytes was shown in Supplementary Figure 1.

### Quantseq

Raw QuantSeq reads were first processed using bbduk (v36.32) ^57^ to remove low quality reads/bases and adapters following the procedure recommended by Lexogen (https://www.lexogen.com/quantseq-data-analysis/). Briefly, kmers were identified based on the Illumina Truseq adapters (included in the BBmap package) and the polyA tail with the following parameters: k=13, ktrim=r, forcetrimleft=11, useshortkmers=t, mink=5, qtrim=t, trimq=10, and minlength=20. The quality-filtered the reads were then mapped to the GRCh28 reference genome (patch12, coding sequences). Because QuantSeq only sequences the 3’ end of the transcripts, read alignment was conducted using bowtie2 (v2.2.8) ^45^ instead of a split-junction aligner. The number of reads mapped to each transcript was then computed using samtools (v1.8) ^53^. Finally, the differentially abundant transcripts were identified using the standard differential expression analysis workflow wrapped in the DESeq function in the R package DESeq2 ^58^.

### Strain-level composition profiling of mWGS samples

The haplotypes and colonization dynamics of the Bacteroides strains were inferred using StrainFinder, which has been successfully applied to track strain dynamics in human-human FMT experiments ^26^. For each Bacteroides species analyzed, metagenomic sequencing reads were first mapped to a reference gene set using bowtie2 (v2.2.8, “very-sensitive” mode) ^45^ to identify SNP sites and extract variant coverage information. We used the species-specific marker genes in the MetaPhlAn2 database ^54^ as the reference gene set to minimize the interference of sequencing reads generated from closely-related species. The resulting sam files were filtered using the StrainFinder pre-processing pipeline with the following parameters: minimum percent identity=90%, minimum alignment length=40, minimum base quality=20, minimum mapping score=0, minimum depth=2, minimum mean coverage=2, maximum acceptable deviation of coverage=2, and default values for the rest of the parameters. Finally, strain haplotypes and relative abundances were inferred by running StrainFinder with the following parameters: dtol=1, ntol=3, max_time=36000 seconds, n_keep=3. The best model – the model with the most probable number of strains (2≤n≤10) in the full dataset – was estimated using model selection based on the Akaike information criterion (AIC).

The compositions of the *A. muciniphila* strains, which were of lower abundances compared to the Bacteroides species, were estimated by assigning sequencing reads to a closest reference strain using Pathoscope 2.0, which was shown to be accurate even at strain resolution ^59,60^. Metagenomic sequencing reads were first mapped to a set of 56 reference genomes downloaded from GenBank (Supplementary Table 3) using Bowtie2 (v2.2.8, “very sensitive” mode) ^45^, followed by a reassignment to the most-likely reference genome using Pathoscope v2.0. The resulting composition profiles of 14 strains (Supplementary Table 3) that had a within-species relative abundance >5% in at least one sample were shown (Figure 5a). The core-genome phylogenetic tree of the 14 strains was constructed using Parsnp (v1.2) ^61^ with a randomly selected reference genome (parameter -r !).

### Association between population states and population features

Strain composition profiles of the microbiome samples – the relative abundances of conspecific strains within each species – were clustered using a Gaussian mixture model implemented in the R package Mclust ^62^. The most likely number of clusters (i.e. Gaussian components) was determined using BIC-based model selection implemented in the Mclust package. Each cluster defines an alternative population state of that species, which was then tested for its association with features of the population – the genotype of the host and the time point of sampling.

The p-values for the association of population states to population features were computed using an unrestricted permutation approach. Given an observation that the majority (X%) of samples sharing a same population feature were classified to only Y population states, we asked whether the association is statistically significant. By definition, p-value is the probability of obtaining an association at least as extreme as the observed simply by chance. To estimate the p-value, the feature of interest was shuffled across all samples, and the frequency of observing at least X% of the feature-bearing samples constrained in at most Y population states (that is, at least as extreme as the real observation) was computed. The final p-value was estimated from 1000 permutations.

### Identification and annotation of A. muciniphila genes

All mWGS samples were pooled and assembled using using MEGAHIT (v1.0.6) ^46,47^ with default parameters. Taxonomic labels of the resulting contigs were predicted using Kraken (v0.10.6) ^63^ to identify *A. muciniphila* contigs. Genes were then predicted from the *A. muciniphila* contigs using Prodigal (v2.6.3) ^64^ and annotated using eggNOG-mapper (v1, database v4.5.1) ^65^. Read-coverages of the *A. muciniphila* genes were estimated by mapping the sequencing reads back to the genes using Bowtie2 (v2.2.8, with parameter -k 10) ^45^, and the number of reads mapped to a given gene was computed using samtools (v1.8) ^53^ with default parameters.

### Statistics

All statistical tests were conducted in R (v3.2.3) using the standard library unless otherwise noted. PERMANOVA was conducted using the adonis function in the vegan package in R. Pairwise PERMANOVA was conducted using the pairwiseAdonis package in R ^66^. Unrestricted permutation was implemented using a custom R script.

The effect of ABT or FMT treatment on engraftment efficiency of human leukocytes in the long-term experiment (Figure 3f) was modeled using a linear model, with treatment, time, and their interaction term compiled using the gls function in the nlme package in R ^67^ while correcting for autocorrelation (using the correlation structure corAR1 implemented in the nlme package. Significance of treatments on the engraftment efficiency was then tested using ANOVA.

To test the enrichment of carbohydrate metabolism and transport genes (Figure 5c), we modeled the null hypothesis assuming that the sampling of gene functions in each gene cluster were Bernoullian, with the success rates (f) given by the overall frequencies of a gene function and the number of trials (n) given by the total number of genes in the group of interest (that is, group III). Therefore, the p-value can be given by the cumulative binomial distribution function:

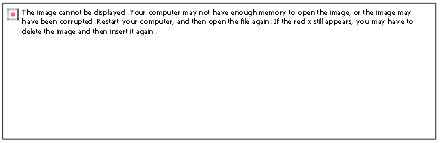

where k is the observed number of genes in the cluster that were annotated with a focal gene function. The p values were computed for all COG gene functional categories present in group III before adjusted via the Benjamini-Hochberg methods.

## Acknowledgment

We would like to thank Ed Keniston, James Keck, John Graham, and Dina’s team for their valuable support in the mouse experiment. We would also like to thank Mark Adams and the JAX microbial genomics core for their support in sequencing, as well as Anthony Carcio and Ted Duffy for their help in immunophenotyping.

## Author contributions

J.O. and K.P. conceived the project. W.Z. and P.K. analyzed the data. J.O. and W.Z. drafted the manuscript. All authors read and approved the final manuscript.

## Figure legends

**Supplementary Figure 1. Identification of different subsets of human leukocytes in a huCD34 mouse**. Analysis of the peripheral blood samples of one mouse collected 0, 14, and 42 days after FMT was shown as an example.

**Supplementary Table 1**. High quality metagenomic bins assembled from the pre-FMT mouse gut samples.

**Supplementary Table 2**. Specifications of the mWGS samples used in this study.

**Supplementary Table 3**. *A. muciniphila* reference genomes used in this study.

**Supplementary Table 4**. Mice used in the long-term immunophenotyping experiments. The peripheral blood samples of mice from the same groups were pooled before downstream analyses.

**Supplementary Table 5. Human panel antibodies used for the immunophenotyping experiments**.

**Supplementary Method 1**. Procedure for preparing cell suspensions from different mouse tissue samples.

