## Supplementary figures and images for "Mice with a humanized immune system are resilient to transplantation of human microbiota"

### Supplementary Figure 1

0 day post-FMT

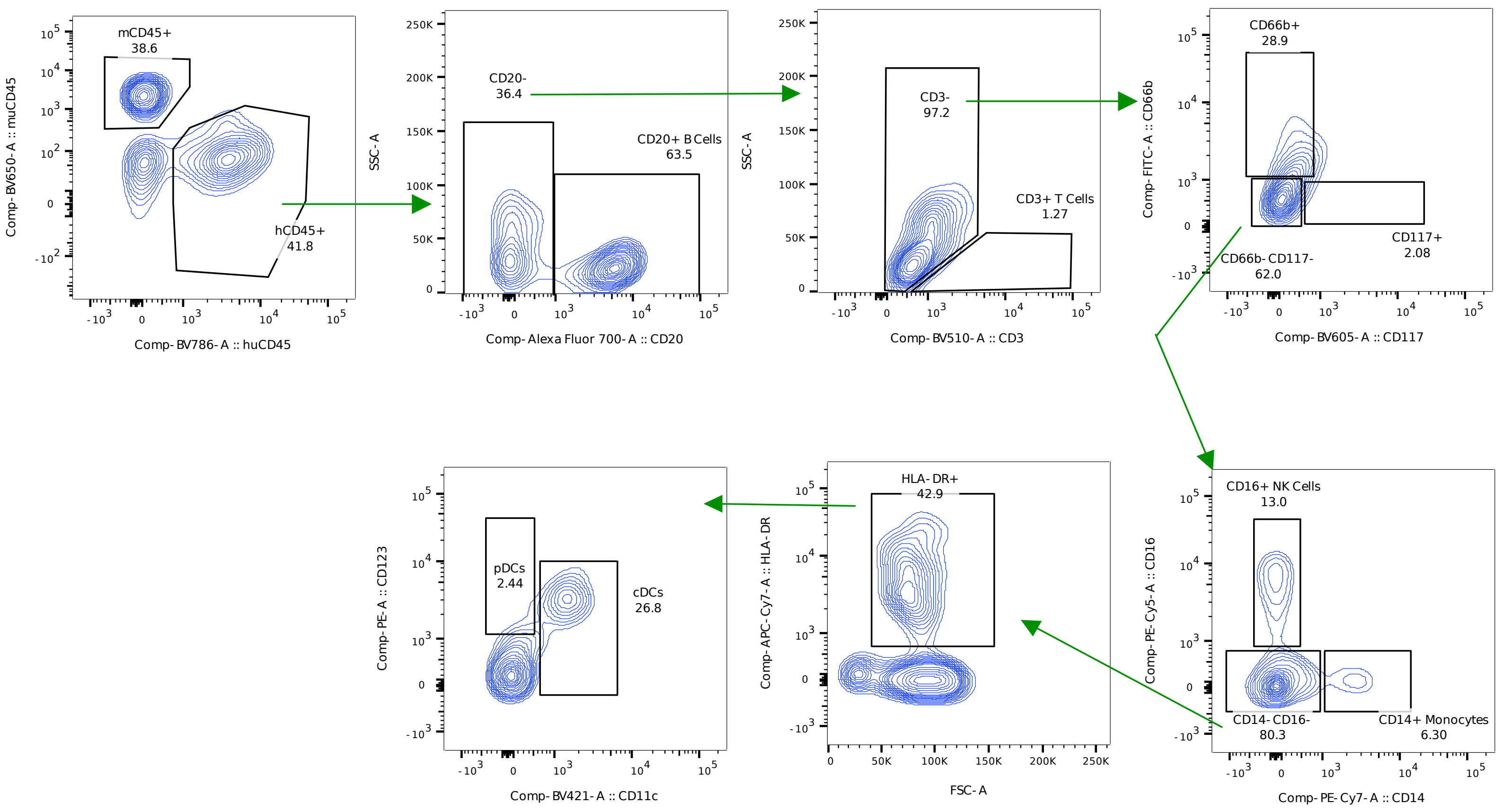


14 days post-FMT

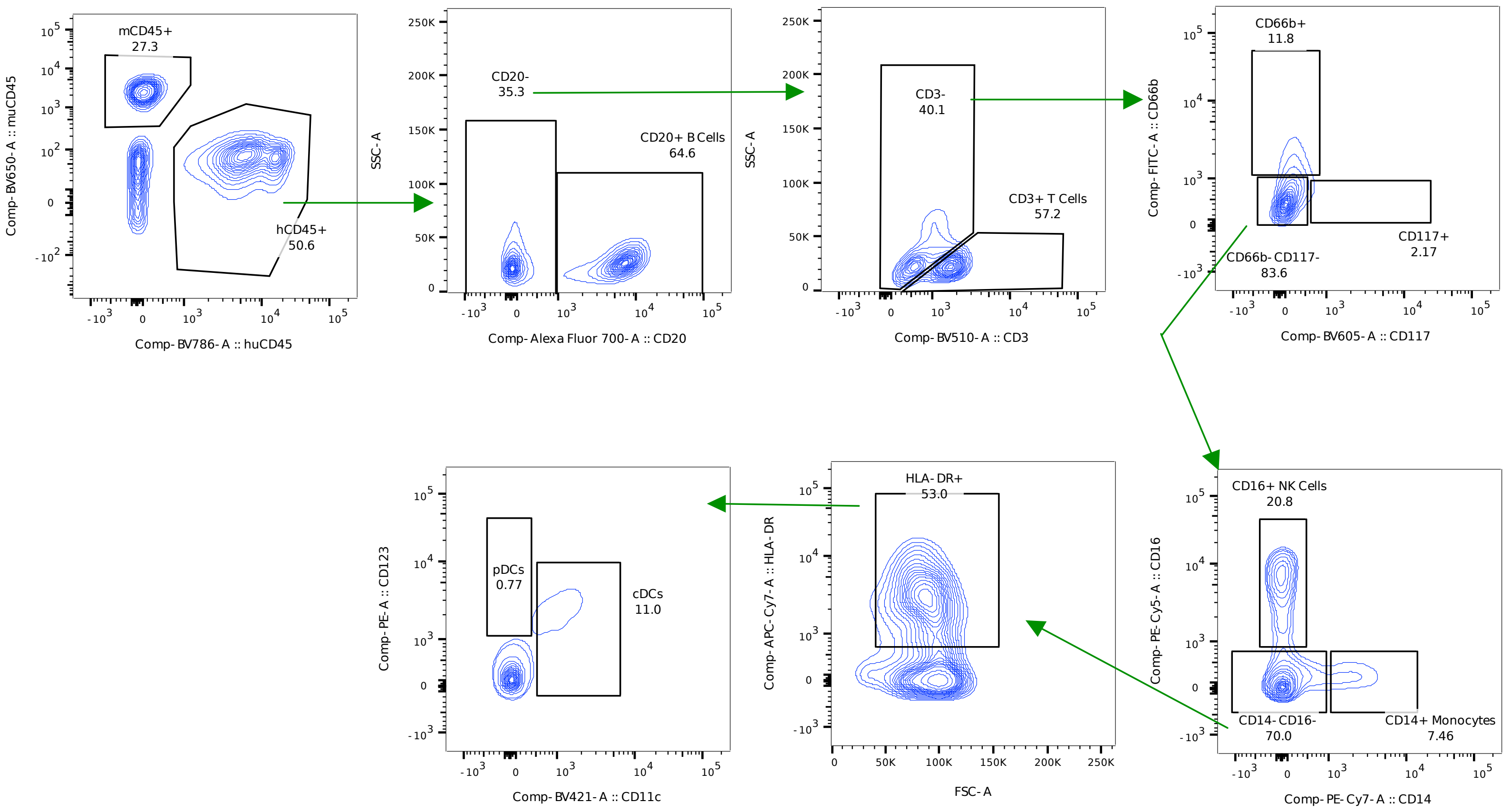


42 days post-FMT

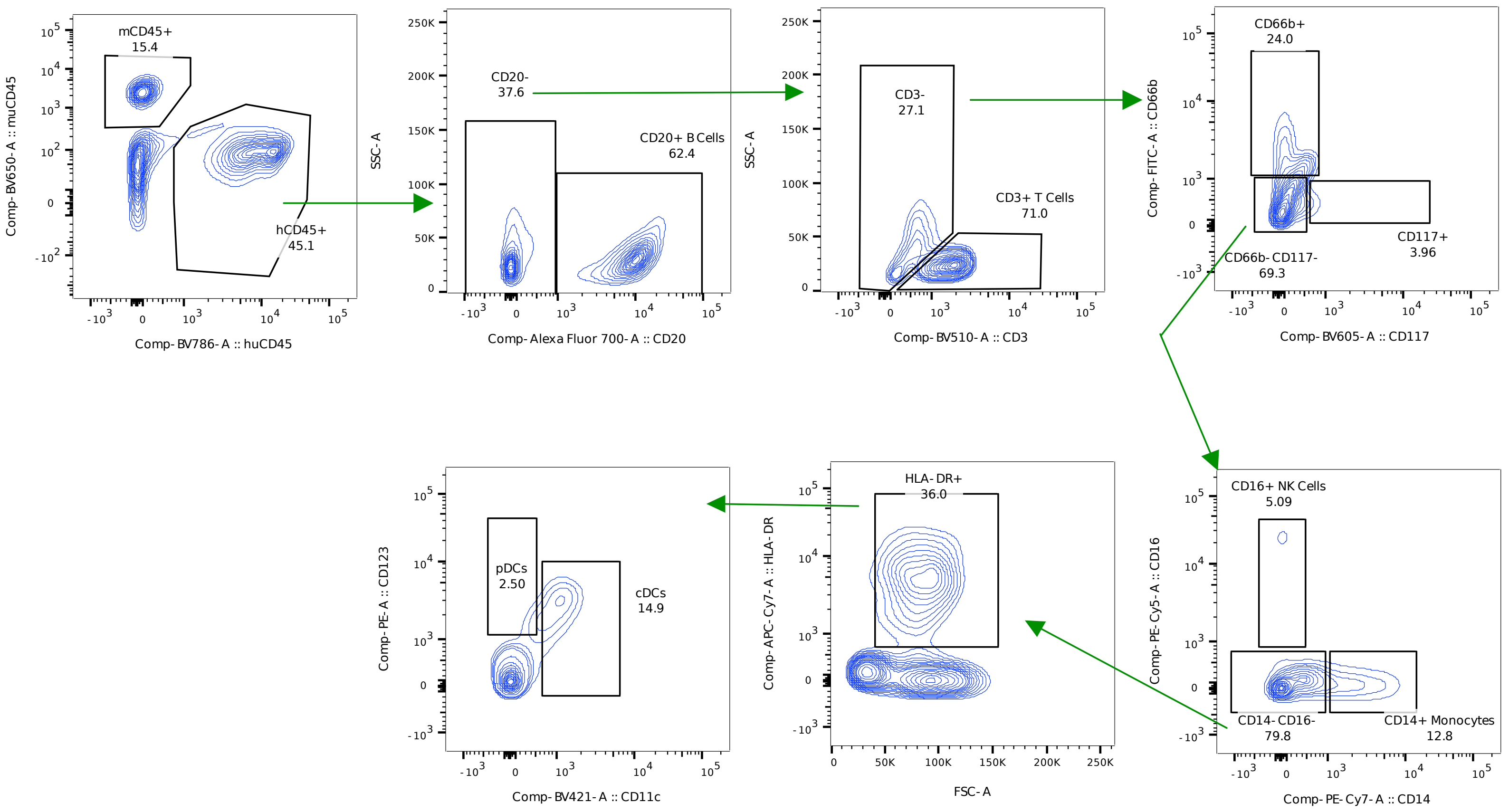
