## Supplementary Method for "Mice with a humanized immune system are resilient to transplantation of human microbiota"

Cell suspension preparation procedures:

Blood—30 min procedure

1. Collect ~1mL of blood with 20ul heparin in 1.5ml tubes.

2. Centrifuge 2500rpm (500xg) for 10 mins and collect the plasma.

3. Freeze plasma at -80°C freezer for future analysis

4. Resuspend cells with 1.5mL RBClysis buffer and incubate for 5 mins

5. Centrifuge 2500rpm 5 mins

6. Remove the supernatant and repeat 4-5 once

7. Resuspend cells with 1mL of PBS-2%FCS

8. Counts the cell number (recover ~50K cells in NSG mice, 500K in Humouse)

9. Resuspend cells in <50M/mL with 10%DMSO/90%FCS and freeze in Freezing box

Bone Marrow—25min procedure

1. Cut out femora and tibiae and remove the muscles with Gauze sponge.

2. Collect bone marrow cells into a petri dish by flushing the shaft with 10 mL of RPMI-10%FCS medium using a 10 mL syringe with a 25-gauge needle.

3. Disaggregate cells by gently pipetting up and down several times using 1mL pipette and transfer into 15mL conical tubes.

4. Centrifuge 1500rpm 5 mins and discard the supernatant.

5. Resuspend cells with 2mL RBClysis buffer and incubate for 5 mins

6. Rinse the cells with 1X PBS and centrifuge 1500rpm 5 mins

7. Resuspend cells with 1mL of PBS-2%FCS and pass the suspension through a 70 μm cell strainer

10. Counts the cell number (recover ~10M cells in NSG mice, 20-30M in Humouse)

11. Resuspend cells in <50M/mL with 10%DMSO/90%FCS and freeze in Freezing box

Spleen—40min procedure

1. Place isolated spleen in a petri dish (6 cm in diameter) containing Liberase and Dnase1 solution to completely cover the bottom of the dish (about 2 mL).

2. Inject mouse spleen with 1mL of enzyme solution per spleen using a 1 mL syringe and a 25G needle

3. Incubate spleen pieces for 10 minutes at 37 °C.

4. Stop the reaction with 5mL of PRMI-10%FBS and make single cell suspension from the spleen using two frosted slides and transfer into 15mL conical tubes.

5. Centrifuge 1500rpm 5 mins and discard the supernatant.

6. Resuspend cells with 2mL RBClysis buffer and incubate for 5 mins

7. Rinse the cells with 1X PBS and centrifuge 1500rpm 5 mins

8. Resuspend cells with 1mL of PBS-2%FCS and pass the suspension through a 70 μm cell strainer

9. Counts the cell number (recover ~10M cells in NSG mice, 20-100M in Humouse)

10. Resuspend cells in <50M/mL with 10%DMSO/90%FCS and freeze in Freezing box
